# Inhibition of MAPK signaling suppresses cytomegalovirus reactivation in CD34^+^ Kasumi-3 cells

**DOI:** 10.1101/2025.02.13.638080

**Authors:** Vargab Baruah, Benjamin A. Krishna, Michael C. Kelly, Xu Qi, Christine M. O’Connor

**Affiliations:** Infection Biology; Sheikha Fatima bint Mubarak Global Center for Pathogen and Human Health Research, Lerner Research Institute, Cleveland Clinic, Cleveland, OH, 44195 USA; Case Comprehensive Cancer Center, Cleveland, OH 44106 USA; Molecular Medicine, Cleveland Clinic Lerner College of Medicine of Case Western Reserve University, Cleveland Clinic, Cleveland, OH 44195 USA

**Keywords:** CMV, latency, MAPK, reactivation, cytomegalovirus, US28

## Abstract

Reactivation of latent human cytomegalovirus (CMV) can lead to severe complications in individuals with dysregulated immune systems. While antiviral therapies for CMV are approved, these compounds are limited by their toxicity and inability to specifically target the latent reservoir or prevent reactivation. Herein we show that CMV reactivation in Kasumi-3 cells, a CD34^+^ hematopoietic cell line, requires mitogen-activated protein kinase (MAPK) activation. Importantly, pharmacological inhibition of the MAPK signaling pathway, including MEK and ERK, restricts viral reactivation in Kasumi-3 cells. In sum, our findings show MAPK signaling is critical for CMV reactivation, revealing a potential avenue for therapeutic intervention to prevent viral reactivation and downstream pathogenesis that is often detrimental for immunosuppressed and immunocompromised patients.

## 1. Introduction

Human cytomegalovirus (CMV) is a prevalent betaherpesvirus that infects a significant portion of the global population. In developed countries, roughly 40–60% of individuals are affected, with rates as high as 100% in underdeveloped regions (Griffiths and Reeves, 2021). Although widespread, primary CMV infections are typically asymptomatic or mildly symptomatic in healthy individuals. Through mechanisms which remain poorly understood, the virus establishes a latent infection in bone marrow CD34^+^ hematopoietic progenitor cells (HPCs), as well as circulating CD14^+^ monocytes. Limited production of infectious virus, coupled with evasion of the host immune response, allows the virus to latently persist within the host for life. Throughout this lifelong infection, latent CMV can sporadically reactivate upon cellular differentiation, leading to lytic production of infectious virus, though these reactivation events are typically asymptomatic in healthy individuals. However, both reactivation and primary infections pose significant risks to those who are immunocompromised, immunosuppressed, or immunonaïve and can lead to severe complications or even death (Goodrum, 2022). For instance, in immunocompromised individuals such as those with AIDS, CMV reactivation frequently results in retinitis, which can ultimately lead to blindness (Springer and Weinberg, 2004). CMV-associated diseases are also more prevalent in solid organ transplant recipients undergoing immunosuppressive regimens, with disease rates as high as 75% in heart-lung transplant recipients (Azevedo et al., 2015). Additionally, primary infection of the immune-naive fetus is the most common congenital infection worldwide, carrying a high risk of damage to the central nervous system, resulting in neurodevelopmental delays, hearing loss, and vision impairment (Akpan and Pillarisetty, 2024).

CMV latent infection is defined by the maintenance of the viral genome, coupled with suppression of viral lytic gene expression and virion production. Many labs have dedicated great effort towards understanding the cellular and viral factors that control the balance between latent and lytic infection. From these studies, the major immediate early (MIE) locus emerges as a key regulator in this process (Goodrum, 2022). The MIE promoter/enhancer, along with the alternative promoters in the MIE locus, drive the expression of the alternatively spliced genes, *UL123* and *UL122*, whose protein products, IE1 and IE2, respectively, transactivate downstream viral loci critical to the success of the viral lytic life cycle (Dooley and O’Connor, 2020). Thus, perhaps it is unsurprising that the MIE locus is highly repressed during latency, allowing for silencing of the viral lytic cascade. Further, many factors are involved in retaining the repression of this locus, including histones and histone remodeling proteins and transcription factors (Dooley and O’Connor, 2020; Goodrum, 2022). Work from a number of groups has revealed recruitment of such factors that promote latency or reactivation is often dictated by altered cellular signaling. Indeed, latent CMV infection reshapes cellular signaling axes, due in part to cmv-miR expression, as well as viral-encoded proteins, such as UL138 and the viral G protein-coupled receptor (GPCR), US28 (Smith et al., 2021). Latent expression of US28 in primary CD14^+^ monocytes or the THP-1 monocytic cell line results in the attenuation of cellular MAPK signaling (Krishna et al., 2017; Wass et al., 2022), leading to a reduction in cellular Fos expression and activity (Krishna et al., 2019), a downstream target of the MAPK pathway (Okazaki and Sagata, 1995). As a component of the transcription factor, Activator Protein-1 (AP-1), the downregulation of Fos in turn prevents the binding of AP-1 to the MIE locus (Krishna et al., 2019). This is a critical step in MIE repression, as AP-1 recruitment to the MIE enhancer/promoter is required for viral reactivation (Krishna et al., 2020a), and is thus, associated with a de-repressed/active MIE locus (Isern et al., 2011). Further, and in line with this, our data showed activation of the MAPK pathway is required for viral reactivation in CD14^+^ primary cells (Wass et al., 2022). Collectively, these findings reveal the importance of MAPK signaling in balancing CMV latency and reactivation, highlighting a potential avenue for novel therapies.

Although antiviral therapies for CMV exist, they have several challenges and limitations. Drugs like ganciclovir and cidofovir are hindered by their toxicity, which restricts both their dosage and effectiveness. Ganciclovir, typically the first-line treatment, is associated with adverse effects such as neutropenia, thrombocytopenia, and nephrotoxicity. Meanwhile, cidofovir, used as a second-line option, also poses risks, notably nephrotoxicity (Bottino et al., 2023). Additionally, these treatments frequently confer viral resistance to multiple drugs, including ganciclovir, letermovir, and maribavir (e.g., refs. (Boonsathorn et al., 2019; Chou et al., 2024; Santos Bravo et al., 2022). In light of these challenges, and in the absence of a vaccine, novel antiviral therapies are imperative. Furthermore, none of the existing therapies specifically target the latent CMV reservoir or downstream reactivation. This necessitates an understanding of the events governing latency and reactivation at the molecular level, which will inform development of new therapeutic strategies. Given how closely MAPK signaling is linked to the regulation of CMV latency and reactivation, exploring novel antiviral interventions targeting this pathway holds promise.

Here, we show that MAPK activity is required for CMV reactivation in Kasumi-3 cells, a CD34^+^ hematopoietic cell line used to model CMV latency (O’Connor and Murphy, 2012). Our findings reveal the pharmacological inhibition of MAPK signaling reverses the lytic-like phenotype in Kasumi-3 cells infected with a US28-deletion mutant. Additionally, targeting MEK or ERK, two key proteins in the MAPK signaling axis, restricts reactivation of wild type CMV in Kasumi-3 cells stimulated with a differentiation/reactivation agent. Collectively, our work highlights the critical role for MAPK signaling in regulating the balance between CMV latency and reactivation and emphasizes the potential of targeting this pathway for therapeutic intervention.

## 2. Materials and Methods

### 2.1. Cells and Viruses

Kasumi-3 cells (ATTC) were maintained in Roswell Park Memorial Institute (RPMI) 1640 medium (ATTC, catalog no. 30-2001), supplemented with 20% fetal bovine serum (FBS), 100 U/ml each of penicillin and streptomycin, and 100 μg/mL gentamicin. Kasumi-3 cells were cultured at a density of 5×10^5^ – 1×10^6^ cells/ml. THP-1-pSLIK-hygro (empty vector) and THP-1-pSLIK-US28-3xF were previously described (Krishna et al., 2019) and cultured in RPMI-1640, supplemented with 10% FBS and 100 U/ml each of penicillin and streptomycin and maintained at a density of 3×10^5^ – 8×10^5^ cells/ml. Primary newborn human foreskin fibroblasts (NuFF-1, passage 13-28; GlobalStem, Rockville, MD, USA) were maintained in Dulbecco’s modified Eagle medium (DMEM), supplemented with 10% FBS, 2 mM L-glutamine, 0.1 mM non-essential amino acids, 10 mM HEPES, and 100 U/ml each of penicillin and streptomycin. All cells were cultured at 37°C and 5% CO_2_.

The bacterial artificial chromosome (BAC)-derived TB40/E*mCherry* (wild type; WT), and TB40/E*mCherry*-US28Δ (US28Δ) viruses were previously described (Miller et al., 2012; O’Connor and Shenk, 2011). All virus stocks were generated as p0 stocks by transfection of BAC DNA into naïve NuFF-1 cells, after which p1 stocks were expanded on naïve NuFF-1 cells, essentially as previously described (O’Connor and Miller, 2021). Virus stocks were then titered on naïve NuFF-1 cells by 50% tissue culture infectious dose assay (TCID_50_).

### 2.2. Other Reagents and Pharmacological Inhibitors

Doxycycline hydrochloride (DOX; ThermoFisher Scientific) was added (1.0 μg/ml) to THP-1-pSLIK-hygro or THP-1-pSLIK-US28-3xF for 24 h, where indicated in the text. DMSO (equivalent v/v) was used as the diluent and vehicle control.

Where indicated in the text and/or Figure Legends, the following pharmacological inhibitors were used (the protein target is indicated in parentheses): U0126 (MEK1/2; ThermoFisher Scientific), selumetinib/AZD6244 (MEK1; Selleck Chemicals), trametinib (MEK1/2; Selleck Chemicals), SCH772984 (ERK1/2; Selleck Chemicals). Optimal concentrations for each inhibitor were determined by the specific protein’s inhibition as well as cell viability (see below). For all pharmacological inhibitors, DMSO (equivalent v/v) was used as the diluent and vehicle control. Specific concentrations and timing of treatment are indicated in the text.

### 2.3. Protein and RNA Analyses

To evaluate protein abundance, total cell lysates were collected at the times indicated in the text and/or Figure Legends and lysed in RIPA buffer (1.0% NP-40; 1.0% sodium deoxycholate; 0.1% SDS; 0.15 M NaCl; 0.01 M NaPO4, pH 7.2; 2 mM EDTA) on ice for 1 hour (h), vortexing every 15 minutes (min). Protein concentrations were determined by Bradford assay with Protein Assay Dye Reagent Concentrate (Bio-Rad). Samples used to determine US28 expression were denatured at 42°C for 10 min. All other samples were denatured at 95°C for 10 min. For all samples, equal protein concentrations were separated by SDS-PAGE and transferred to nitrocellulose by semi-dry transfer. The following antibodies were used at the indicated concentrations: anti-FLAG clone M2 (MilliporeSigma, 1:7,500), anti-phospho-p44/42 MAPK (ERK1/2; Thr202/Tyr204) (Cell Signaling Technology [CST], 1:1,000), anti-p44/42 MAPK (ERK1/2; CST, 1:1,000), anti-beta-actin-peroxidase (MilliporeSigma, 1:20,000), and goat-anti-mouse horseradish peroxidase (HRP) secondary (Jackson ImmunoResearch Labs, 1:10,000).

To quantify cellular transcripts, total RNA was harvested and then purified using the High Pure RNA Isolation kit (Roche) in accordance to the manufacturer’s protocol. Equal RNA concentrations (1.0 μg) were then used to generate cDNA with random hexamer primers and TaqMan Reverse Transcription (RT) Reagents (ThermoFisher Scientific), and equal volumes of cDNA were then used for quantitative polymerase chain reaction (qPCR). The following primers were used: Fos, forward – 5’-CACTCCAAGCGGAGACAGAC-3’ and reverse – 5’-AGGTCATCAGGGATCTTGCAG-3’; HMGCR, forward – 5’-TGATTGACCTTTCCAGAGCAAG-3’ and reverse – 5’-CTAAAATTGCCATTCCACGAGC-3’. All samples were analyzed by qPCR in triplicate using a 96-well-format CFX Connect Real Time PCR machine (Bio-Rad).

### 2.4. Toxicity Assays

NuFF-1 (1.75 x 10^4^) or Kasumi-3 (2 x 10^4^) cells were seeded onto 96-well opaque white CulturPlates (Perkin Elmer). NuFF-1 cells were allowed to adhere overnight. For NuFF-1 cells, cultures were incubated with inhibitors for 24 h, after which cells were washed three times with 1X PBS. NuFF-1 cell viability was evaluated 2 and 5 d later (i.e., 3 and 6 d post-treatment) to allow for sufficient cell doubling. Kasumi-3 cells were treated with inhibitors for 24 and 48 h, at which point cell viability was assessed. For all pharmacological inhibitors, DMSO (equivalent v/v) was used as the diluent and vehicle control. Puromycin (20 µM to 0.08 µM) and hygromycin (15 mM to 0.06 µM) were used as positive controls in NuFF-1 and Kasumi-3 cells, respectively. For each compound, a series of nine two-fold dilutions were used, spanning the following concentration ranges: U0126, 0.625 – 160 µM; SCH772984, 0.008 – 2 µM; selumetinib, 0.03 – 8.0 µM; trametinib, 0.006 – 1.6 µM. Cell viability was assessed by CellTiter-Glo (Promega) according to the manufacturer’s instructions, and plates were read on a Cytation 5 (Agilent) plate reader.

### 2.5. Latency and Reactivation Assays

Kasumi-3 cells were cultured in XVIVO-15 (Lonza) for 2 days (d) prior to infection. Cells were then infected with the indicated viruses at a multiplicity of 1.0 TCID_50_/cell, as previously described (O’Connor and Murphy, 2012). In brief, the infection of the Kasumi-3 cells was centrifugally enhanced (1000 x *g*, 35 min, room temperature) at 5×10^5^ cells/ml in XVIVO-15. The infected cells were then cultured overnight, after which they were treated with trypsin to remove virus that did not undergo cell entry. Infected cultures were then cushioned onto Ficoll-Paque (GE Healthcare Life Sciences), washed three times with phosphate buffered saline (PBS), replated at 5×10^5^ cells/ml in XVIVO-15, and returned to culture. Infected cells were again cushioned onto Ficoll-Paque at 5 days post-infection (dpi) and replated at a density of 5×10^5^ cells/ml in XVIVO-15. Kasumi-3 cells were treated with the indicated pharmacological inhibitor or equivalent volume/volume (v/v) dimethylsulfoxide (DMSO; ThermoFisher Scientific) at the concentrations and time noted in the text. Where indicated, infected Kasumi-3 cells were treated with 20 nM 12-O-Tetradecanoylphorbol-13-acetate (ThermoFisher Scientific) or equivalent volume/volume (v/v) DMSO (ThermoFisher Scientific) for 2 d to differentiate the cells and reactivate latent virus. Latently-infected or differentiated Kasumi-3 cells were then washed with 1X PBS and co-cultured with naïve NuFF-1 cells in the absence of compounds to quantify the frequency of infectious centers by Extreme Limiting Dilution Analysis (ELDA; (Hu and Smyth, 2009), http://bioinf.wehi.edu.au/software/elda/), as previously detailed (Peppenelli et al., 2021).

### 2.6. Quantification of viral lytic replication

NuFF-1 cells were pretreated with each inhibitor at the indicated concentrations for 1 h. Cells were then infected with TB40/E*mCherry* (moi = 1.0 TCID_50_/cell) for 1 h with gentle agitation every 15 min. Infected cells were then washed three times with 1X PBS, and fresh media with inhibitors were replaced for 24 h. For all pharmacological inhibitors, DMSO (equivalent v/v) was used as the diluent and vehicle control. Inhibitor- or DMSO-containing media was then removed, cells were washed as before with 1X PBS, and fresh media (without inhibitor or DMSO) was replenished. At 96 hpi, media was collected for TCID_50_ assay using naïve NuFF-1 cells.

## 3. Results

### 3.1. U0126 treatment suppresses infectious virus production in US28Δ-infected Kasumi-3 cells

We and others have shown US28 is required to maintain a latent infection in myeloid cells (Miller and O’Connor, 2024). Indeed, infection of myeloid cells with a virus lacking the US28 ORF (TB40/E*mCherry*-US28Δ; US28Δ) fails to establish latency and results in the production of infectious virus (Humby and O’Connor, 2015; Krishna et al., 2019; Mahmud et al., 2024; Wass et al., 2022). Additionally, in monocytes, TB40/E*mCherry* (wild type; WT) latent infection results in an attenuation of MAPK signaling via US28, though activation of this pathway is required for reactivation (Wass et al., 2022). Whether fine-tuning of this signaling pathway is critical to latency and reactivation in CD34^+^ cells, however, remained unclear. Given our finidngs in monocytes and the requirement for US28 to maintain latency in both primary monocytes and CD34^+^ cells, we reasoned these two cell types might share the need to attenuate MAPK signaling to maintain latency. To this end, we first evaluated the efficacy of the potent MEK1/2 inhibitor, U0126, as MEK1/2 is a protein that is upstream in the MAPK signaling axis. Our assessment of U0126 toxicity in Kasumi-3 cells, a CD34^+^ hematopoietic cell line that serves as a model system for CMV latency (O’Connor and Murphy, 2012), revealed minimal impact on cell health at 24 h, though some toxicity at higher concentrations at 48 h (**Fig. 1A**). U0126 treatment for 24 h was effective at inhibiting ERK activity (**Fig. 1B**). Due to the minimal toxicity we observed at 24 h, we validated our assay using hygromycin as a control (**Fig. S1A**), which confirmed our toxicity assay was functioning as intended at the 24 h time point. Next, we tested the impact of restricting MEK1/2 activity on Kasumi-3 cell infection. To this end, we infected Kasumi-3 cells with WT or US28Δ under latent conditions for 7 days (d), after which we treated the infected cultures with U0126 or vehicle (DMSO, v/v) for an additional 2 d at 10 μM, which has no impact on cell viability (**Fig. 1A**). We then washed the cells and evaluated the ability of these cultures to maintain latency using extreme limiting dilution analysis (ELDA) by co-culturing the infected populations with naïve fibroblasts. Our data reveal inhibiting MEK1/2 with U0126 does not impact latency in WT-infected cells (**Fig. 2** and **Fig. S2**, blue bars). However, and supporting our prior work (Humby and O’Connor, 2015), US28Δ-infected Kasumi-3 cells fail to maintain a latent infection (**Fig. 2** and **Fig. S2**, open green bar), although suppression of MEK1/2 via U0126 treatment restricts the lytic-like phenotype we observe with the US28Δ-infected Kasumi-3 cells (**Fig. 2** and **Fig. S2**, green checked vs. open bars). Additionally, this U0126-mediated effect we observed is not due to suppression of lytic infection or cell health of the fibroblasts. We performed parallel experiments, where we treated fibroblasts with U0126 or DMSO (v/v) for 24 hours (h), after which we washed the cells free of compound, and replenished them with fresh media. We allowed the cells to undergo two rounds of doubling, assessing their viability at 3 and 6 d post-treatment. We found U0126 treatment of fibroblasts reveal little-to-no cell toxicity (**Fig. 3A**). Indeed, our positive control using puromycin treatment of fibroblasts displayed cellular toxicity at higher doses, as expected (**Fig. S1B**). Additionally, we tested the effect of U0126 treatment on lytic replication of virus in fibroblasts. To this end, we pre-treated fibroblasts with various concentrations of U0126 or DMSO (v/v) for 24 h. We then washed the cells to remove the compounds and infected the cells with WT, and following adsorption, washed the cells and replenished the cultures with fresh media containing U0126 or DMSO (v/v) at their original concentrations. After 24 h, the U0126- or DMSO-containing media was removed, cells washed, and fresh media replenished. We then returned the cells to culture and collected the media at 96 hours post-infection (hpi) to quantify viral titers by 50% tissue culture infectious dose (TCID_50_). Our findings reveal U0126 treatment also results in a decrease in lytic viral growth, though the differences were not statistically significant (**Fig. 3B**). As there is a reduction in titer during lytic infection, perhaps it is unsurprising that there is also an impact on reactivation; indeed it is likely that the activity of some pathways in these phases of infection are shared. In sum, these data suggest inhibiting the MAPK axis via MEK1/2 allows the US28Δ-infected Kasumi-3 cells to retain their latent state.

**Figure 1.**
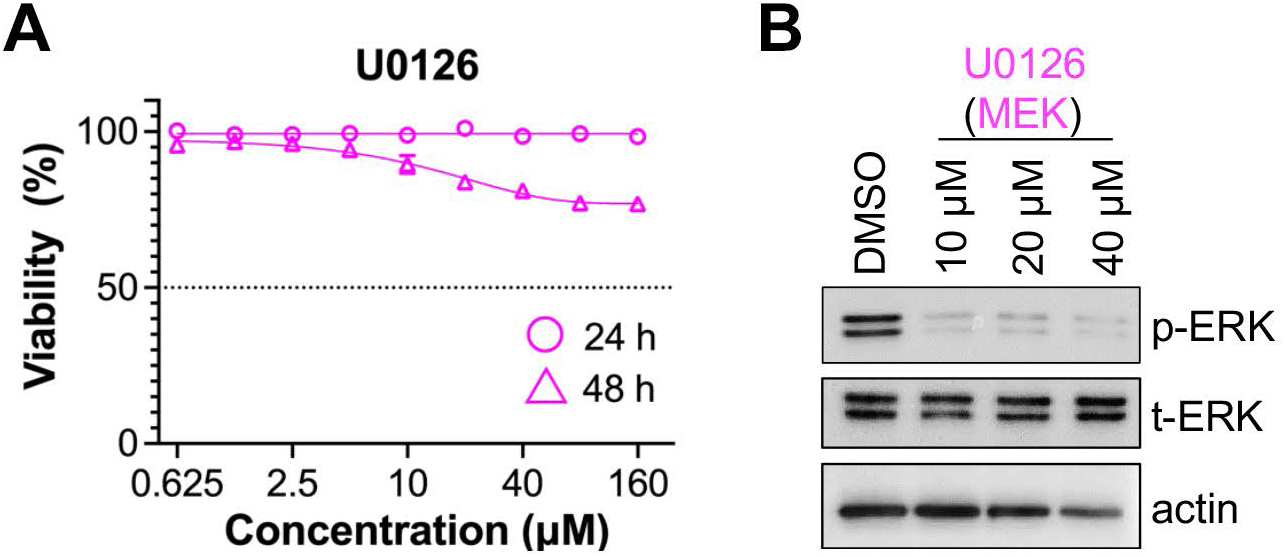
The MEK1/2 inhibitor, U0126, inhibits downstream ERK activity in Kasumi-3 cells with minimal toxicity at 24 h post-treatment. (A) Kasumi-3 cells were either untreated (DMSO) or treated with U0126 for 24 or 48 h. Cell survival was evaluated in the absence (DMSO, v/v) or presence of U0126 at varying concentrations. Percent cell viability is shown relative to cells treated with vehicle (DMSO), assessed at 24 h and 48 h post-treatment. Hygromycin is shown in Figure S1A as a positive control. N=3, representative assay shown. (B) Kasumi-3 cells were treated with DMSO (v/v) or U0126 at the indicated concentrations for 24 h. Cultures were harvested and lysates probed for the indicated proteins by immunoblot. N=3, representative blots shown.

**Figure 2.**
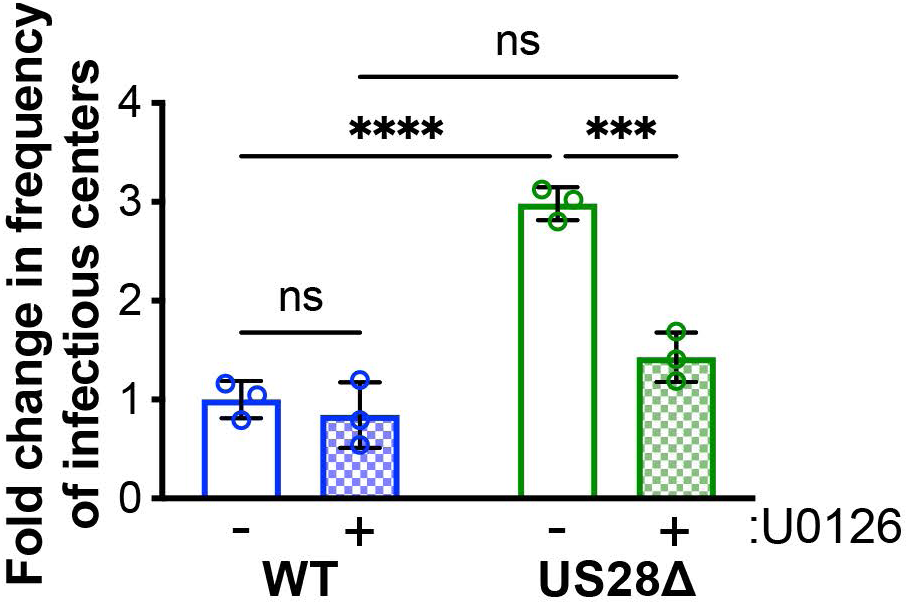
Treatment with the MEK1/2 inhibitor, U0126, limits lytic-like replication in Kasumi-3 cells. Kasumi-3 cells were infected (moi = 1.0 TCID_50_/cell) with WT or US28Δ under latent conditions. At 7 dpi, half of each culture was treated with DMSO (-) or 10 μM U0126 (+) for 2 d, after which the cells were washed and co-cultured with naïve fibroblasts to measure the fold change in frequency of infectious centers relative to WT (-U0126, open blue bar) by ELDA. Each data point (circles) denotes the mean of three technical replicates. Error bars indicate the standard deviation (SD) of the three biological replicates (**Figure S3**). Statistical significance was calculated by two-way ANOVA with Tukey’s test for multiple comparisons. \*\*\**p*<0.001, ns - not significant

**Figure 3.**
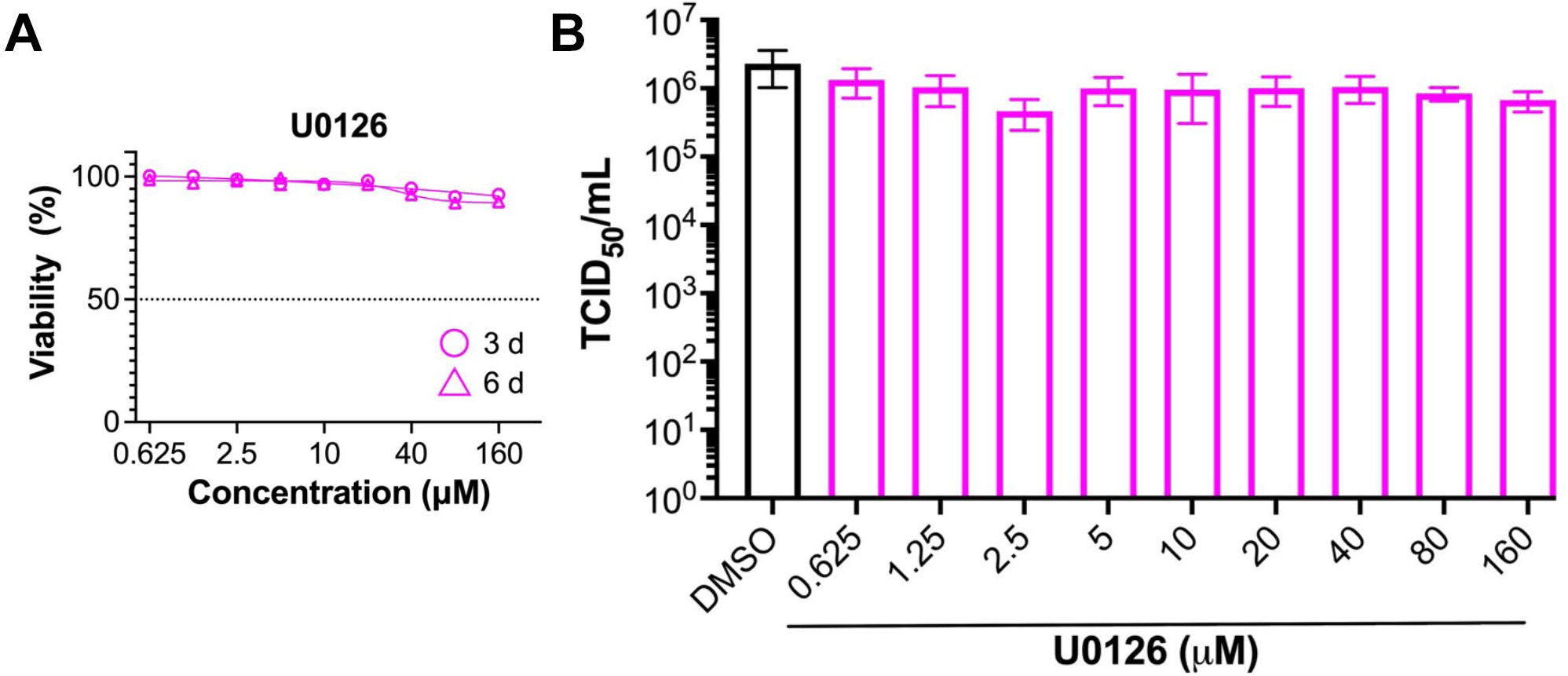
U0126 treatment results in minimal toxicity in primary fibroblasts and does not significantly impact viral lytic growth. (**A**) NuFF-1 fibroblasts were treated with U0126 for 24 h, then washed with PBS and returned to culture. Percent cell viability is shown relative to cells treated with vehicle (DMSO), assessed at 3 and 6 d post-treatment. Puromycin is shown in **Figure S1B** as a positive control. N=3, representative assay shown. (**B**) NuFF-1 cells were pretreated for 24 h with DMSO (v/v/) or U0126 at the indicated concentrations. Cells were washed with PBS and infected with TB40/E*mCherry* (moi = 1 TCID50/cell) for 1 h. Viral inoculum was removed, and cells were washed with 1X PBS. Media containing DMSO (v/v) or U0126 at the indicated concentrations was then replenished. Following 24 h in culture, DMSO- or drug-containing media was removed, cells were washed, and fresh media (without U0126 or DMSO) was replenished. Cells were returned to culture, and media was collected at 96 hpi. Viral titers were quantified by TCID50 assay on naïve NuFF-1 fibroblasts. N=3 biological replicates, each with N=3 technical replicates; representative assay shown.

### 3.2 MEK1/2 inhibition or US28 expression attenuates FOS expression and protein abundance

Using other latency model systems, we previously showed US28 downregulates *FOS* transcript levels, which functions to aid in the maintenance of viral latency (Krishna et al., 2019). Since MAPK signaling regulates Fos (Okazaki and Sagata, 1995), we hypothesized US28-mediated *FOS* attenuation may result from upstream downregulation of MAPK signaling via US28. To this end, we utilized our previously generated THP-1 monocytic cells in which we overexpressed US28 under the control of a doxycycline (DOX)-inducible promoter (Krishna et al., 2019). Upon DOX-treatment of US28 expressing cells, we observed a robust attenuation of *FOS* mRNA expression relative to control cells (**Fig. 4A**), consistent with our prior work (Krishna et al., 2019). Furthermore, US28-mediated suppression of *FOS* transcription is similar to that quantified in U0126-treated THP-1 cells, regardless of US28 expression (**Fig 4A**), suggesting inhibition of MEK1/2 activity reduces *FOS* expression. To test whether U0126 treatment impacted US28-mediated protein levels, we treated control and US28-expressing cells with DMSO (v/v) or DOX for 24 h in the presence U0126 or DMSO (v/v). U0126 treatment alone results in attenuated phospho-ERK, phospho-Fos, and total Fos (**Fig. 4B**), suggesting the inhibition of MEK1/2 impacts this pathway in these cells. Furthermore, DOX-induced expression of US28 results in attenuation of both the phosphorylated and total forms of Fos, as well as repression of phosphorylated ERK (**Fig. 4B**). While the impact of US28 expression alone on phospho-ERK is not as great as U0126 treatment, US28 expression does result in a significant decrease in the phosphorylated form of this MAPK pathway protein. It is important to note that DOX treatment results in a significant upregulation of phospho- and total Fos, which we and others have noted previously (Fujioka et al., 2004; Krishna et al., 2019; Toton et al., 2012). However, DOX-induced expression of US28 indeed attenuates both forms, as does U0126 treatment (**Fig. 4B**). Collectively, these data suggest MEK1/2 inhibition or US28 expression tempers the activity of MAPK signaling.

**Figure 4.**
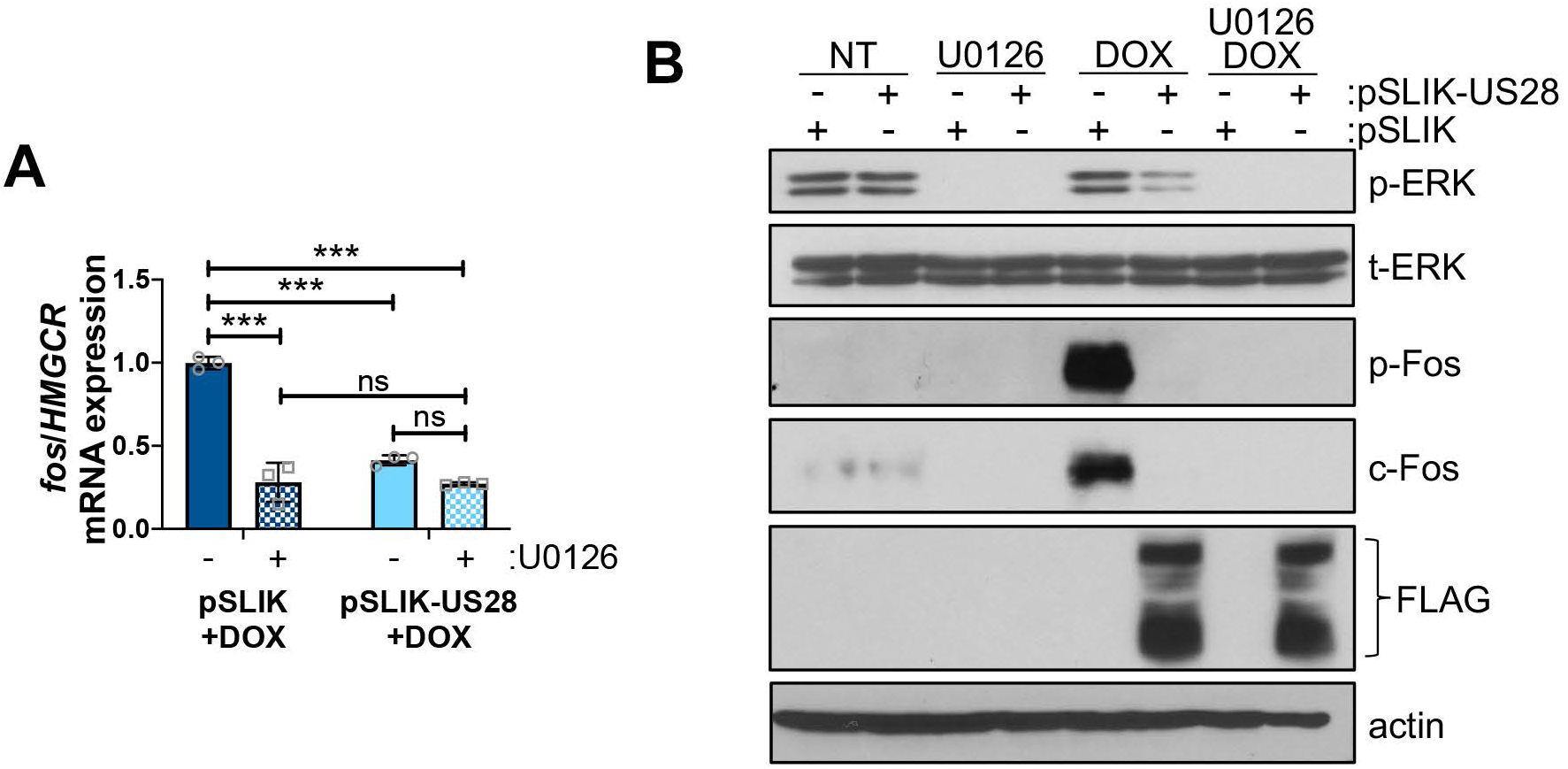
Fos is attenuated by U0126 treatment or US28 expression. (**A**) THP-1-pSLIK-hygro (empty vector control; pSLIK) or THP-1-pSLIK-US28-3xF (pSLIK-US28) were treated with DOX (24h) in the absence (-; DMSO, v/v) or presence (+) of 10 μM U0126. *FOS* gene expression was measured relative to cellular *HMGCR*. *** *p* < 0.001; ns, not significant. Representative experiment shown; N = 2 (**B**) THP-1 cells in (**A**) were treated with DMSO (v/v) or DOX in the absence (DMSO, v/v) or presence of 10 μM U0126 for 24h. Each cell type was cultured in the absence of either compounds (not treated; NT) and instead treated with DMSO (v/v). Lysates were analyzed by immunoblot for phospho- or total-ERK (p-ERK, t-ERK), p-Fos, c-Fos, US28-3xF (with a FLAG antibody), or actin. Representative blots shown; N = 3.

### 3.3. The MAPK signaling axis is important for CMV reactivation in Kasumi-3 cells

Given our above findings, including that U0126 reversed the lytic-like phenotype we usually observe in US28Δ-infected Kasumi-3 cells, we hypothesized increased MAPK activation is critical for viral reactivation in this CD34^+^ cell line. To test this, we utilized more targeted pharmacological inhibitors for the MAPK pathway, including SCH772984 (ERK1/2), selumetinib (MEK1), and trametinib (MEK1/2). Similar to our work with U0126, we first evaluated toxicity and potency of these compounds in Kasumi-3 cells. We treated Kasumi-3 cells with each compound or vehicle (DMSO, v/v) and evaluated cell viability at 24 and 48 h. Our data reveal very minimal toxicity for SCH772984 (**Fig. 5A**), selumetinib (**Fig. 5B**), or trametinib (**Fig. 5C**), compared to our control treatment of hygromycin (**Fig. S1**). Additionally, we evaluated the potency of each compound and found 125 nM SCH772984, 0.5 µM selumetinib, and 0.1 µM trametinib effectively inhibited ERK1/2 phosphorylation in Kasumi-3 cells (**Fig. 5D,E**). Thus, to test whether MAPK inhibition restricts CMV reactivation, we infected Kasumi-3 cells with WT or US28Δ under latent conditions for 7 d. We then differentiated the cell populations with TPA for an additional 2 d, thereby allowing viral reactivation, in the presence or absence of each inhibitor using concentrations that attenuate protein activity, while not impacting overall cell health (**Fig. 5**). We next evaluated the ability of the infected cultures to maintain latency and support reactivation using ELDA, as above. We found inhibiting ERK1/2 (125 nM SCH772984) or MEK (0.5 µM selumetinib, 0.1 µM trametinib) results in the reversal of the US28Δ-infected cell ‘lytic-like’ phenotype, even in the absence of TPA treatment (**Fig. 6**, **Fig S3**), similar to our observations following U0126 treatment (**Fig. 2** and **Fig. S2**). Importantly, each compound restricts WT reactivation (**Fig. 6**, **Fig S3**), suggesting robust activation of the MAPK signaling axis is important for viral reactivation. Similar to our work with the U0126 inhibitor, these results are not likely due to the inhibitors impacting fibroblast health or their ability to support infection. We performed similar control experiments for SCH772984, selumetinib, and trametinib as we did for U0126 (**Fig. 3**). We found minimal fibroblast toxicity associated with SCH772984, selumetinib, or trametinib at the 3 d time point (**Fig. 7**) when compared to our control treatment with puromycin (**Fig. S1B**), although SCH772984 treatment resulted in toxicity at higher concentrations. Each compound resulted in increased toxicity at higher concentrations at the 6 d timepoint (**Fig. 7**). We observed statistically significant effects on viral titers following compound treatment (∼2-3 fold) (**Fig. 7B,D,F**). It is also important to note that in our latency/reactivation assays, we are treating the Kasumi-3 cells with the inhibitor compounds and washing the cells free of compounds or vehicle prior to co-culturing these cells with the fibroblasts. Thus, it is unlikely the fibroblasts are impacted by the inhibitors (or vehicle) in our ELDA assays. This perhaps is unsurprising, as pathways required for successful reactivation from latency are likely important for robust lytic infection. In sum, our findings reveal the attenuation of the MAPK signaling axis is critical for maintaing viral latency, though its activation is required for efficient CMV reactivation. Taken together, these data suggest targeting the MAPK pathway effectively suppresses CMV reactivation in Kasumi-3 cells.

**Figure 5.**
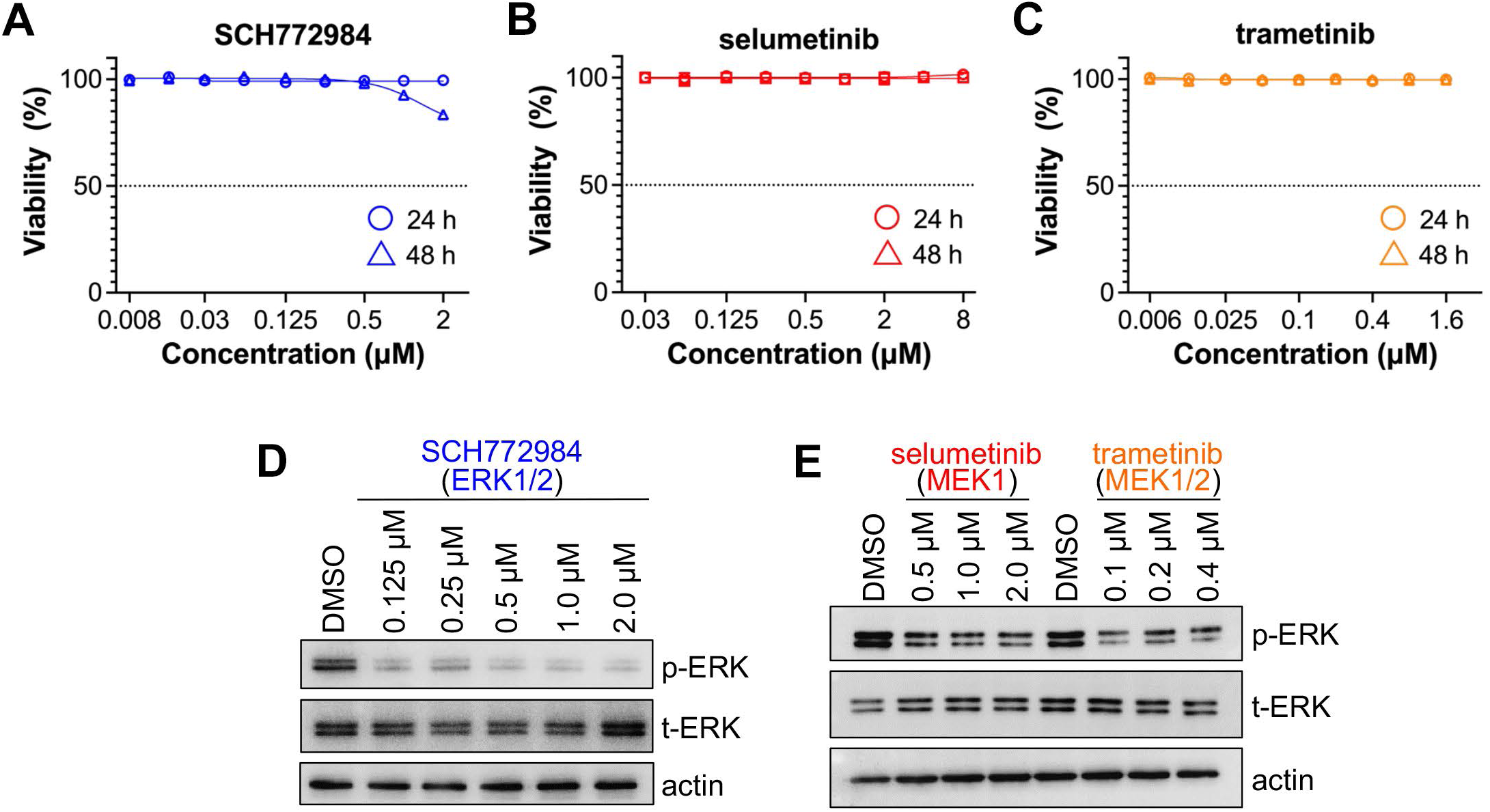
MAPK inhibitors attenuate ERK phosphorylation. Kasumi-3 cells were treated with DMSO (v/v) or (**A,D**) SCH772984, (**B,E**) selumetinib, or (**C,E**) trametinib at increasing concentrations. (**A-C**) Cell viability was assessed at 24 and 48 h; percent cell viability is shown relative to cells treated with vehicle (DMSO), assessed at 24 h or 48 h post-treatment. Hygromycin is shown in **Figure S1A** as a positive control. N=3, representative assays shown for each inhibitor. (**D,E**) Cells were treated for 24 h and lysates were harvested for immunoblot and probed for the indicated proteins. N=3, representative blots shown.

**Figure 6.**
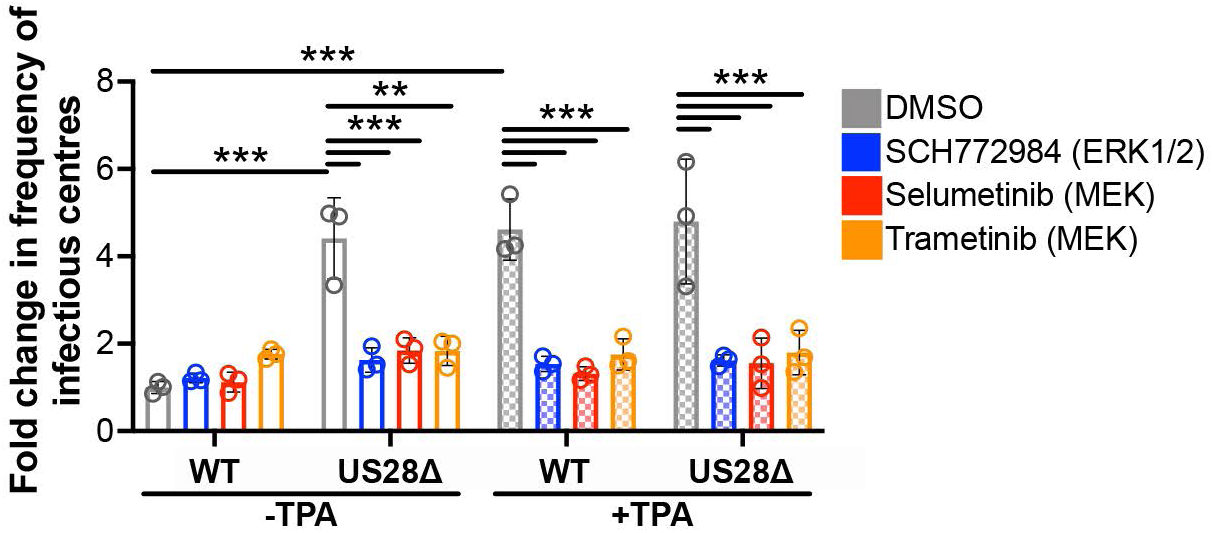
Inhibiting MEK/ERK attenuates viral reactivation in Kasumi-3 cells. Kasumi-3 cells were infected (moi = 1.0 TCID_50_/cell) with WT or US28Δ for 7 d and then treated with the indicated compounds for an additional 2 d in the presence of DMSO (-TPA) to maintain latent conditions or TPA (+TPA) to differentiate cells, allowing for reactivation. Cultures were washed to remove the compounds, and the frequency of infectious centers was quantified by co-culturing each cell population with naïve fibroblasts using ELDA software (Hu and Smyth, 2009), http://bioinf.wehi.edu.au/software/elda/. Data is plotted as fold-change in frequency of infectious centers relative to untreated (DMSO, -TPA), WT-infected cells (open gray bar). Each data point (circles) denotes the mean of three technical replicates. Error bars indicate the SD of the three biological replicates (each shown in **Figure S3**). Concentrations for the compounds are as follows: 125 nM SCH772984, 0.5 µM selumetinib, 0.1 µM trametinib). Statistical significance was calculated by two-way ANOVA with Tukey’s test for multiple comparisons. \*\**p* < 0.05, \*\*\**p* < 0.001.

**Figure 7.**
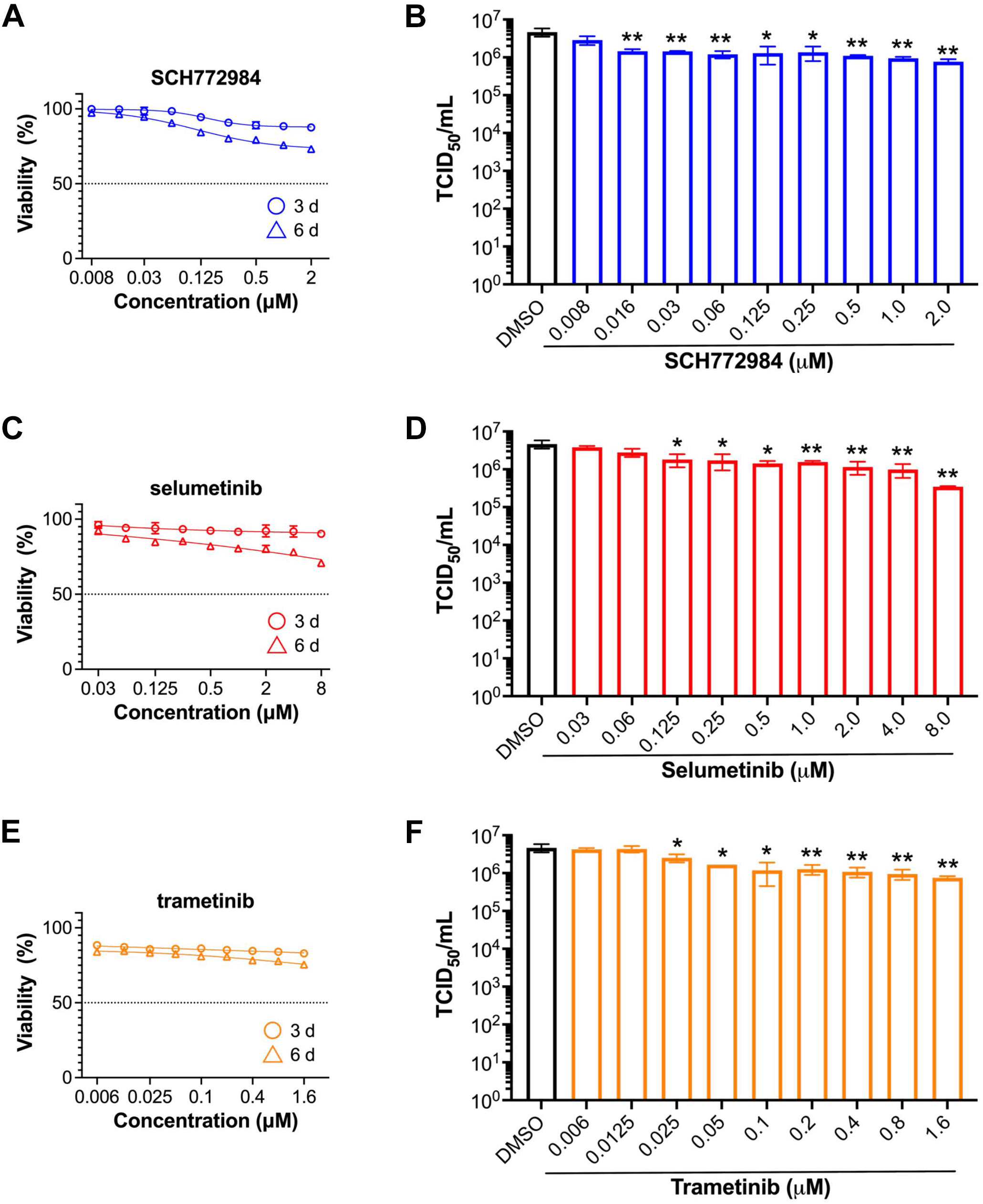
MEK/ERK inhibitors display minimal toxicity and impact on viral growth in fibroblasts. (**A, C, E**) NuFF-1 fibroblasts were treated DMSO (v/v) or the indicated compound at increasing concentrations for 24 h, washed, and fresh media replenished. Cell viability was determined at 3 or 6 d post-treatment and percent viability is shown relative to cells exposed to vehicle. Puromycin is shown in Figure S1B as a positive control. N=3, representative assay shown for each compound. (**B, D, F**) NuFF-1 cells were pretreated for 24 h with DMSO (v/v/) or the pharmacological compound at increasing concentrations, as indicated. Cells were washed with 1X PBS and infected with TB40/E*mCherry* (moi = 1.0 TCID_50_/cell) for 1 h. Viral inoculum was removed, cells were washed with 1X PBS, and media containing DMSO (v/v) or the compound at the indicated concentrations was then replenished. Following 24 h in culture, DMSO- or drug-containing media was removed, cells were washed, and fresh media (without compound or DMSO) was replenished. Cells were returned to culture, and media was collected at 96 hpi. Viral titers were quantified by TCID_50_ assay on naïve NuFF-1 fibroblasts. Note the DMSO control was used as the control for each inhibitor treatment, as the v/v of DMSO required was the same. N=3 biological replicates, each with N=3 technical replicates; representative assay shown. \**p*<0.05; \*\**p*<0.01

## 4. Discussion

A number of labs have demonstrated the diverse and pivotal roles of MAPK signaling in CMV infection. For example, Kew et al. provided insights into the multifaceted response triggered by CMV through ERK-MAPK signaling to promote the survival of latently infected cells (Kew et al., 2017). Additionally, CMV regulates latency and lytic replication states by modulating epidermal growth factor receptor (EGFR) trafficking (Buehler et al., 2016), a cellular receptor that activates MAPK. We recently showed that while CMV infection results in activation of EGFR upon virion entry into primary CD14^+^ monocytes, US28 tempers this cellular receptor’s activity, which skews downstream signaling (Mahmud et al., 2024). Indeed, our work in primary CD14^+^ monocytes also revealed the critical, concerted roles of US28-mediated MAPK signaling in maintaining viral latency in CD14^+^ monocytes (Wass et al., 2022). More recently, building upon these prior findings, our current data demonstrate the necessity of this signaling axis for viral reactivation in the less differentiated, CD34^+^ Kasumi-3 hematopoietic cell line. Our data herein reveal latently-infected Kasumi-3 cells stimulated with TPA to differentiate the cells and reactivate virus fail to reactivate to WT levels when treated with MAPK inhibitors. Specifically, inhibiting the MAPK pathway with a variety of compounds targeting MEK (selumetinib, trametinib) and ERK (SCH772984) effectively prevents WT reactivation in latently-infected Kasumi-3 cells. Furthermore, treating US28Δ-infected Kasumi-3 cells with MEK (selumetinib, trametinib, U0126) or ERK (SCH772984) inhibitors attenuated the lytic-like phenotype typically observed with this virus, in-line with our previous work in CD14^+^ monocytes (Wass et al., 2022). Collectively, our findings further highlight the importance of MAPK signaling in regulating the balance between viral latency and reactivation, revealing a potential pathway to target to prevent viral reactivation.

Despite our findings, it is important to note Buehler and colleagues demonstrated inhibition of MEK (binimetinib) or ERK1/2 (SCH772984) enhances viral reactivation in primary, cord blood-derived CD34^+^ HPCs (Buehler et al., 2019). These observations are likely explained by the intrinsic differences between these cell types. While Kasumi-3 cells are a viable model for CMV latency (O’Connor and Murphy, 2012), there are undoubtedly differences between the transformed cell line and primary, cord blood-derived CD34^+^ HPCs cultured *ex vivo*. For instance, while Kasumi-3 cells express CD34, these cells display characteristics of undifferentiated leukemia cells and notably, display high levels of *EVI1* expression (Asou et al., 1996). Further, Kasumi-3 cells also express CD33 and CD38 (Albright and Kalejta, 2013; Asou et al., 1996), suggesting these cells are further differentiated than the primary CD34^+^/CD38^-^ HPCs used in latency studies (Peppenelli et al., 2021). Therefore, while Kasumi-3 cells faithfully recapitulate many aspects of latency and reactivation when compared to primary CD34^+^ HPCs (e.g., refs. (Albright and Kalejta, 2013; Humby and O’Connor, 2015; Kim et al., 2019; Krishna et al., 2019; Krishna et al., 2021; Krishna et al., 2022; Krishna et al., 2020a; Krishna et al., 2020b; O’Connor and Murphy, 2012; O’Connor et al., 2016; O’Connor et al., 2014; Sandhu and Buchkovich, 2020; Schwartz et al., 2023), infections in the two latency models are likely not identical. This is not unprecedented, as latently-infected Kasumi-3 cells are non-responsive to the HDAC inhibitor, valproic acid, whose treatment does lead to reactivation in primary CD34^+^ HPCs (Albright and Kalejta, 2013). Nonetheless, our current and prior (Wass et al., 2022) findings align with previous work in CD14^+^ monocytes differentiated to immature interstitial cell-like dendritic cells, in which MAPK activity promoted CMV reactivation (Reeves and Compton, 2011). Reeves and Compton showed differentiation/reactivation was driven via inflammation-associated signaling triggered by IL-6. MAPK signaling activated mitogen and stress-activated kinases, resulting in the phosphorylation of the transcription factor, CREB, and histone H3 at serine 10, collectively promoting a lytic environment (Reeves and Compton, 2011). This finding also underscores the capacity of MAPK signaling to influence CMV through multiple triggers and effectors.

Our data also reveal CMV modulates cellular MAPK signaling in Kasumi-3 cells, at least in part through the viral GPCR, US28, to balance the latent and lytic phases of infection. At which levels of MAPK signaling might US28 exert its effect? We previously showed US28 upregulates and interacts with the host receptor tyrosine kinases, Ephrin receptor A2 (EphA2), influencing signaling at an early, upstream level of the MAPK pathway (Wass et al., 2022). In addition, because US28 is incorporated into the mature CMV particle (Humby and O’Connor, 2015), its immediate availability in an infected cell enables CMV to rapidly modulate host signaling, as we have recently demonstrated (Mahmud et al., 2024). However, due to the intricacies of MAPK signaling, it is certain there are additional complexities to consider, especially as the MAPK pathway interfaces with other signaling axes to serve common cellular functions. Indeed, several other MAPK-adjacent signaling pathways, involving Wnt, Akt, and phospholipase C are implicated in CMV latency and reactivation (de Wit et al., 2016; Langemeijer et al., 2012; Miller et al., 2012). CMV also engages with the host cell via EGFR, potentially intertwining with EphA2-mediated signaling, and regulating a shared subset of effectors (Fulkerson et al., 2020; Kim et al., 2016). Buehler et al. demonstrated roles for the CMV proteins, pUL135 and pUL138, in regulating host EGFR. While pUL135 promotes the internalization and turnover of surface EGFR to drive lytic replication and reactivation, pUL138 preserves EGFR’s surface expression and activation during latent infection of CD34^+^ HPCs, thereby promoting latency (Buehler et al., 2016). Furthermore, host EGFR signaling creates positive feedback on pUL138 by inducing its expression during latency via MAPK signaling. However, our own work in primary CD14^+^ monocytes reveal US28 also tempers EGFR activity upon viral entry. This results in skewed Akt activity, which aids in the establishment of viral latency (Mahmud et al., 2024). Indeed, and as mentioned above, Buehler and colleagues found that inhibiting the MEK/ERK pathway downstream of EGFR increases viral reactivation in latently infected CD34^+^ HPCs (Buehler et al., 2019). Therefore, it is possible MAPK signaling is differentially regulated in CD34^+^ versus CD14^+^ cells or the cell line we tested (Kasumi-3 cells). It is also important to note that signaling is not binary; in fact, subtle changes in pathway activities result in profound changes in the cell environment, which may also be cell type specific. In line with this, CMV activates MAPK signaling to counter pro-apoptotic factors such as Bak, promoting transcription of key survival factors and ensuring the survival of infected myeloid progenitor cells as a latent reservoir (Kew et al., 2017). Thus, these data suggest US28 does not abolish ERK activity, but rather represses ERK activity to levels conducive for latent infection that must be overcome for successful reactivation. Collectively, these data indicate even subtle differences in protein activity can have profound influence on the outcome of infection. These findings further highlight the fine-tuning of cellular pathways that regulate the delicate balance between CMV latency and reactivation.

Finally, MAPK emerges as a targetable pathway for more than CMV infections; several herpesvirus subfamily members depend on MAPK signaling activation to steer towards lytic behavior. For instance, the activation of MAPK-interacting kinase 1/2 (Mnk) is associated with Epstein-Barr virus (EBV) lytic replication in epithelial cells. Pharmacological inhibition of Mnk using CGP57380 prevents phosphorylation of the Mnk target, eIF4E, consequently inhibiting lytic replication in a dose-dependent manner (Adamson et al., 2022). Kaposi’s sarcoma-associated herpesvirus (KSHV) employs its viral protein kinase, vPK, to phosphorylate ERK and mediate its translocation to the nucleus for subsequent signaling during reactivation and lytic replication (Kim et al., 2020). Infection with herpes simplex virus type 1 (HSV-1) modifies ERK1/2 compartmentalization during active replication, and pharmacological inhibition using U0126 prevents the ERK-dependent cyclin E activity and reduces viral replication and yield (Colao et al., 2017). Even in cases of co-infection with HSV-1 and KSHV, HSV-1-induced ERK/MAPK signaling activity leads to KSHV reactivation (Qin et al., 2011). Thus, targeting the MAPK signaling axis could represent an avenue to target multiple herpesviruses. Many inhibitors targeting MAPK-driven diseases have received FDA approval, including two utilized in this study - selumetinib and trametinib. Selumetinib is approved to treat neurofibromatosis type I, a rare genetic disorder in children (Gross et al., 2020), while trametinib is primarily used in anti-cancer regimens for melanomas and gliomas (Flaherty et al., 2012). In their original intended application, both drugs aim to inhibit MAPK signaling, consequently inhibiting cell proliferation and inducing apoptosis in tumor cells. A similar mechanism is envisioned for CMV therapy, where virus reactivation is attenuated due to depleted MAPK signaling. Importantly, unlike the currently-approved CMV antivirals, toxicity with these MAPK inhibitors is not a concern. Selumetinib’s safety profile, coupled with the absence of cumulative toxic effects, reveal its suitability for long-term treatment (Sharawat et al., 2022). This would prove particularly beneficial for persistent, latent CMV infections. Similarly, trametinib has a favorable pharmacokinetic profile, characterized by a prolonged half-life and minimal toxicity (Infante et al., 2012). Standing FDA approval adds to their appeal, potentially paving a smoother path towards repurposed clinical applications for CMV and perhaps other herpesviruses, as discussed above. Vargas et al. have highlighted such repurposing potential or MAPK pathway inhibitors in an HIV-1 latency model (Vargas et al., 2019). Further investigations into additional components of the MAPK signaling axis, potentially influenced by US28, will prove crucial for developing effective and viable therapies. Such studies will inform downstream clinical trials to assess inhibitor efficacy and safety in preventing viral reactivation in latent CMV carriers, as well as others harboring latent herpesvirus infections.

## 5. Conclusions

Overall, our findings demonstrate MAPK signaling is critical for balancing CMV latency and reactivation. Inhibition of this signaling axis restricts viral reactivation, highlighting a potential target for therapeutic intervention. Furthermore, and consistent with our prior work in primary CD14^+^ monocytes, CMV-encoded US28 modulates this signaling pathway. As a viral GPCR, this is an attractive target for anti-viral therapy, and future work aimed at gaining a complete understanding of how this viral signaling protein impacts latency and reactivation will undoubtedly inform intervention strategies.

## 6. Glossary

BAC: bacterial artificial chromosome
CMV: cytomegalovirus
DMSO: dimethylsulfoxide
DOX: doxycycline
d: days
dpi: days post-infection
ELDA: extreme limiting dilution analysis
ERK: extracellular signal-regulated kinase
GPCR: G protein-coupled receptor
h: hours
HPC: hematopoietic progenitor cells
IE: immediate early
min: minutes
MAPK: mitogen activated protein kinase
MEK: mitogen activated protein kinase kinase
MIE: major immediate early
MIEP: major immediate early promoter
MOI: multiplicity of infection
NuFF-1: primary newborn human foreskin fibroblasts
TCID_50_: median tissue culture infectious dose
TPA: 12-O-tetradecanoylphorbol-13-acetate
UL: unique long
US: unique short

## Acknowledgements

The authors thank Nancy Wang for technical assistance with the toxicity assays and helpful discussions.

