## Supplemental File for "Inhibition of MAPK signaling suppresses cytomegalovirus reactivation in CD34^+^ Kasumi-3 cells"

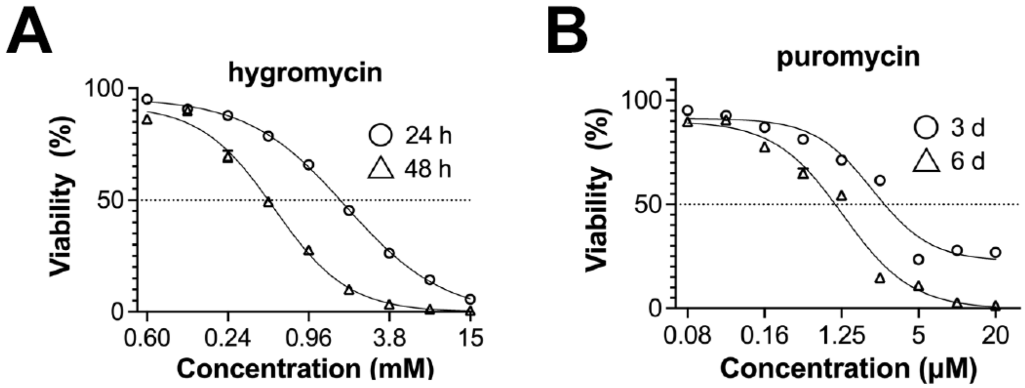


**Figure S1. Kasumi-3 and NuFF-1 fibroblasts display decreased viability at high concentrations of hygromycin and puromycin, respectively.** (**A**) Kasumi-3 cells were either untreated (DMSO, v/v) or treated with hygromycin at increasing concentrations for 24 or 48 h. (**B**) NuFF-1 fibroblasts were treated DMSO (v/v) or puromycin at increasing concentrations for 24 h, washed, and fresh media replenished. Cell viability was determined at 3 or 6 d post-treatment. (**A,B**) Percent cell viability is shown relative to cells treated with vehicle (DMSO), assessed at either (**A**) 24h and 48 h post-treatment or (**B**) 3 and 6 d post-treatment. N=3, representative assays are shown for each compound.

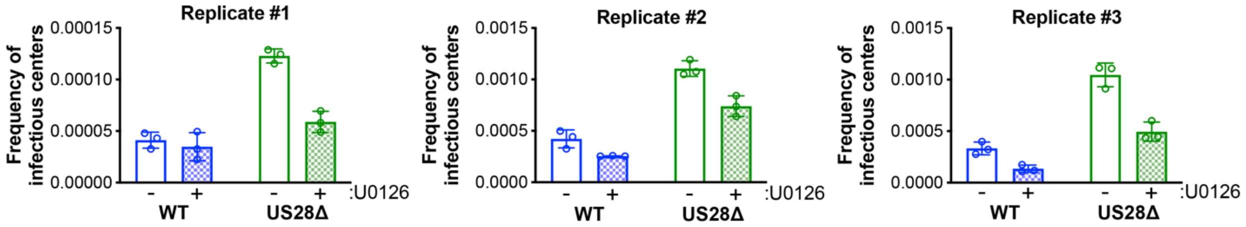


**Figure S2. Individual replicates for the data in Figure 2.** Kasumi-3 cells were infected as described in Figure 2. Infected Kasumi-3 cells were then co-cultured with naïve NuFF-1 cells to measure the frequency of infectious centers by ELDA. Three biological replicates are shown. Each data point (circles) denotes a technical replicate within each experiment.

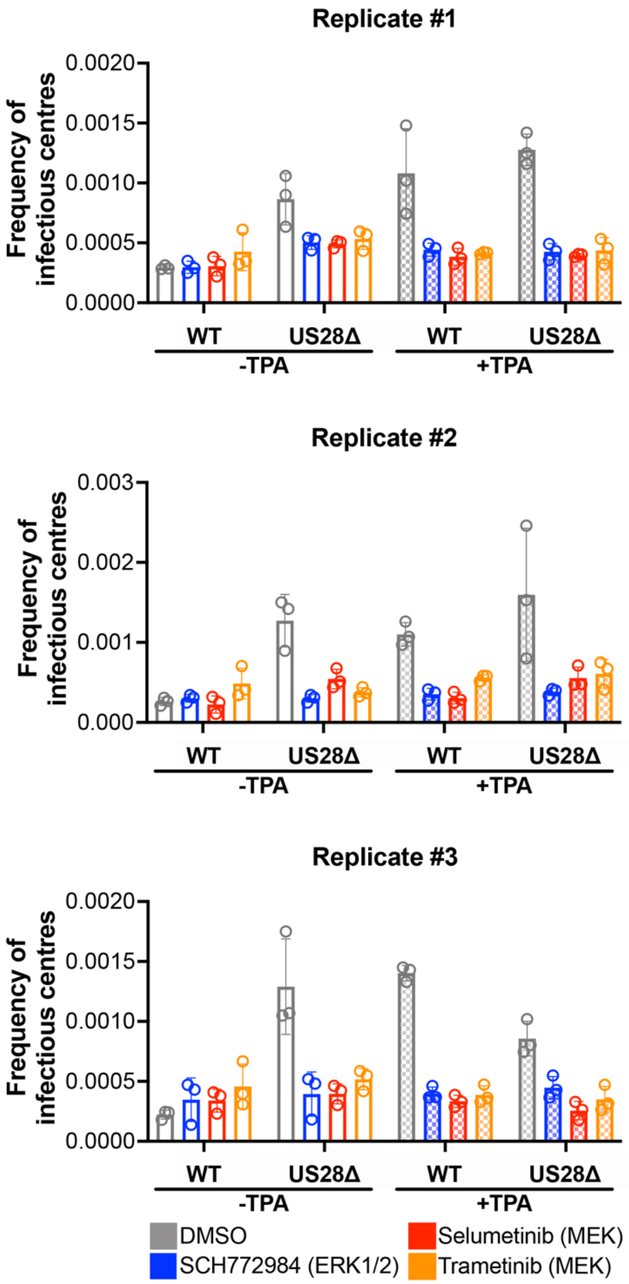


**Figure S3. Individual replicates for the data in Figure 6.** Kasumi-3 cells were infected as described in Figure 6. Infected Kasumi-3 cells were then co-cultured with naïve NuFF-1 cells to quantify the frequency of infectious centers by ELDA. Three biological replicates are shown. Each data point (circles) denotes a technical replicate within each experiment.
